# POmAb, an antibody targeting open prothrombin, results in anticoagulation without excessive bleeding in mice

**DOI:** 10.1101/2025.06.29.662207

**Authors:** Marisa A. Brake, Suresh Kumar, Glenn Merrill-Skoloff, Sol Schulman, Robert Flaumenhaft, Nicola Pozzi

## Abstract

Anti-prothrombin antibodies are commonly found in patients with Antiphospholipid Syndrome (APS), yet their role in clinical manifestations remains unclear. We recently identified two classes of anti-prothrombin antibodies based on their ability to recognize closed and open forms of prothrombin. Type-I antibodies bind to the open form, while Type-II antibodies bind to both forms. POmAb is a prototypical Type-I antibody that specifically targets kringle-1 of prothrombin, maintaining it in an open state. In this study, we assess the effects of POmAb in mice using the cremaster arteriole laser-induced injury model. POmAb bound mouse prothrombin and decreased thrombin generation in mouse plasma. When administered intravenously shortly before the injury, POmAb quickly accumulated on the damaged vessel wall. This accumulation significantly reduced fibrin generation with a modest effect on platelet accumulation and without causing excessive bleeding. Results obtained with POmAb offer insights into the potential roles of the anti-prothrombin antibodies in APS. They also provide proof of concept for a new class of anticoagulants that, by specifically targeting open prothrombin, could mitigate thrombosis with reduced bleeding risk.

## Introduction

Prothrombin is the target of anti-prothrombin antiphospholipid antibodies in the systemic autoimmune disorder Antiphospholipid Syndrome (APS)^1^. Understanding how autoantibodies interact with prothrombin and contribute to the development of APS is crucial for advancing diagnosis and clinical management^2-5^.

Prothrombin exists in closed and open forms, with the closed form being the predominant one in plasma under basal conditions^6-8^. Anti-prothrombin antibodies discriminate between these two forms^9^, with Type-I antibodies preferring the open form and Type-II antibodies binding to both forms^4,9^. Recently, we identified POmAb (<u>P</u>rothrombin <u>O</u>pen <u>m</u>onoclonal <u>a</u>nti<u>b</u>ody) as a prototypical Type-I antibody^10^. Biophysical and cryo-EM structural analyses demonstrated that POmAb specifically binds kringle-1 of prothrombin through an extensive interface centered on the region R90-Y93, resulting in stabilization of the open form and *in vitro* reduction of thrombin generation via a novel mechanism of action^10,11^. How this *in vitro* reduction of thrombin generation due to conformational modulation translates *in vivo* is unknown and is investigated in this study.

## Materials and Methods

### Antibodies and other reagents

POmAb was expressed in CHO cells and purified as described previously^10^. Purity was >95% (4–12% SDS-PAGE under non-reducing conditions) and 99% (SEC-HPLC). Endotoxin level was < 0.1 EU/mg. POmAb concentration was calculated using the following molecular weight and extinction coefficient (E_280nm_^0.1%^): 150,000 Da, 1.58. *In vivo* grade recombinant mouse IgG_1_ isotype control antibody was from Syd Labs (PA007126). Antibodies were aliquoted and stored at -80°C until used. Fluorescently labeled POmAb was prepared by conjugating 1.3 mg of antibody in 0.5 mL of 50 mM sodium borate buffer (pH 8.5) with DyLight 650-NHS ester. Protein/dye ratio was 3.27 moles of dye per mole of protein. Anti-fibrin antibody was purified from a growing hybridoma clone 59D8. Anti-CD42b beta antibody for staining platelets was from Emfret Analytics. All other reagents were purchased from Invitrogen (Carlsbad, CA) and Sigma (St. Louis, MO) unless otherwise stated.

### Enzyme-linked immunoassay

One hundred microliters of 1 μg/ml human, mouse, rat or bovine prothrombin (Prolytix) solubilized in 0.1 M sodium bicarbonate pH 9.6 were added to a Nunc MAXISORP plate and incubated overnight at 4°C. After washing three times with 200 μl/well of 20 mM Tris-HCl, pH 7.4, 145 mM NaCl, 5 mM CaCl_2,_ Tween 20 0.05% (TBSC-T), wells were blocked with 200 μl of 20 mM Tris-HCl pH 7.4, 145 mM NaCl, 1% BSA (TBS-B) for 60 min at room temperature. One hundred microliters of POmAb (0-100 μg/ml) prepared in the sample buffer 20 mM Tris-HCl pH 7.4, 145 mM NaCl, 5 mM CaCl_2,_ Tween 20 0.05%, 1% BSA (TBSC-BT) were added to each well and incubated for 60 min at room temperature. Plates were washed three times TBSC-T, and then 100 μl of 1:10,000 dilution of HRP-conjugated antimouse IgG (γ-chain specific) antibody (Sigma Aldrich) was added for 60 min at room temperature. Plates were washed three times with TBS-T and then incubated with 100 μl of 3,30,5,50-tetramethylbenzidine (TMB) liquid substrate (Sigma Aldrich). After 30 min, the colorimetric reaction was quenched with 100 μl of TMB-stop solution. The optical density at 450 nm was recorded using a SPARK microplate reader (TECAN).

### Competition Experiments

One hundred microliters of 1 μg/ml mouse or human prothrombin (Prolytix) solubilized in 0.1 M sodium bicarbonate pH 9.6 were added to a Nunc MAXISORP plate and incubated overnight at 4°C. After washing three times with 200 μl/well of 20 mM Tris-HCl, pH 7.4, 145 mM NaCl, 5 mM CaCl_2,_ Tween 20 0.05% (TBSC-T), wells were blocked with 200 μl of 20 mM Tris-HCl pH 7.4, 145 mM NaCl, 1% BSA (TBS-B) for 60 min at room temperature. One hundred microliters of 10 μg/ml POmAb (mouse) or 0.1 μg/ml POmAb (human) mixed with 0-20 μM mouse or 0-5 μM human prothrombin in the absence or presence of 400 μM argatroban were prepared in the sample buffer 20 mM Tris-HCl pH 7.4, 145 mM NaCl, 5 mM CaCl_2,_ Tween 20 0.05%, 1% BSA (TBSC-BT), added to each well and incubated for 60 min at room temperature. Plates were washed three times TBSC-T, and then 100 μl of 1:10,000 dilution of HRP-conjugated anti-mouse IgG (γ-chain specific) antibody (Sigma Aldrich) was added for 60 min at room temperature. Plates were washed three times with TBS-T and then incubated with 100 μl of 3,30,5,50-tetramethylbenzidine (TMB) liquid substrate (Sigma Aldrich). After 30 min, the colorimetric reaction was quenched with 100 μl of TMB-stop solution. The optical density at 450 nm was recorded using a SPARK microplate reader (TECAN).

### Diluted Russell’s Viper Venom (dRVV)

dRVV was measured using the HEMOCLOT LA-S (ACK090K-RUO) kit on an ST4 semiautomated coagulometer (Diagnostica Stago, Gennevilliers, France). Briefly, 50 μl of mouse citrated plasma was mixed with 50 μl of antibodies (0-5 μM) in the appropriate cuvette. After 180 seconds at 37°C, the reaction was started by adding 50 μl of dRVV reagent.

### Activation of Prothrombin

Prothrombin activation was monitored using a colorimetric assay that continuously reports the amount of thrombin that is generated upon cleavage by the prothrombinase complex^8,10,12^. Briefly, prothrombin (250 μl, 25 nM) bound to IgG POmAb (0-2.4 μM) was reacted with 2.5 pM factor Xa (Prolytix), 20 μM phospholipids (75:25 POPC: POPS) (Avanti), 0.2 nM factor Va (Prolytix), and 24 μM chromogenic substrate FPF-pNA. Data were collected on a SPARK microplate reader (TECAN).

### Laser-injury model

Thrombus formation in response to laser injury was measured as described previously^13-15^. Briefly, cremaster muscle arterioles were injured using an ABLATE! pulsed laser system (Intelligent Imaging Innovations, Den. CO). Platelet, fibrin, and POmAb accumulation were measured by infusion of Dylight 650-labeled POmAb (0.5 mg/kg body weight), Dylight 405-labeled anti-platelet (CD42b; 0.1 µg/g body weight; Emfret Analytics) and DyLight 488-labeled anti-fibrin (clone 59D8; 0.3 µg/g) antibodies through a jugular vein catheter. Data were acquired before and after laser injury using the brightfield, 370/445 nm, 480/510 nm, and 640/670 nm channels. Images were captured for 250 seconds at 2 frames/s using a CCD camera (ORCA Flash 4.0, Hamamatsu Photonics, Japan) on an Axio Observer fluorescence microscope (Carl Zeiss, Germany). Data were analyzed using Slidebook 6.0 (Intelligent Imaging Innovations). Data from 40 to 48 thrombi were used to determine the median value of the integrated fluorescence intensity to account for variability in thrombus formation under any given experimental condition. AUC was calculated for individual thrombi and normalized to injury lengths to evaluate statistical significance. Injury lengths were determined as previously described^15^.

### Tail bleeding assay

C57BL/6J mice (stock number 000664) were purchased from the Jackson Laboratory. Male and female 9-week-old mice were anesthetized by intraperitoneal injection of 100 mg/kg ketamine and 10 mg/kg xylazine. Mice were infused via internal jugular vein catheter with either 0.5 mg/Kg IgG or POmAb. The tail was transected at a 1.0-mm diameter and placed in 0.9% saline solution as previously described^16^. The total bleed time, number of rebleeds, and hemoglobin (absorbance at 575-nm) was measured. A standard curve generated using pooled C57BL/6J mouse blood was used to convert absorbance to blood volume.

### Rigor and statistical Analysis

Biochemical assays were repeated at least three times using two lots of POmAb preparations. Unless otherwise specified, data were plotted and analyzed using Prism 10.5 with appropriate statistical tests specified in each figure legend.

## Results

Human and mouse prothrombin kringle-1 share 85% sequence identity, with most POmAb-targeted residues—including R90–Y93—conserved **(Figure 1A)**. Accordingly, ELISA showed nanomolar EC_50_ binding of POmAb to mouse prothrombin **(Figure 1B)**. However, this value was ∼50-fold lower than that of human prothrombin. Of 12 non-conserved residues **(Figure 1C)**, only H76, E85, and Q110 interact with POmAb based on the cryo-EM structure of POmAb bound to human prothrombin^10^ **(Figure 1D)**. These are mutated in mouse prothrombin to T, Q, and K, likely reducing binding. Rat and bovine prothrombin also carry mutations at H76 and Q110 but retain E85 **(Figure S1A)** and showed stronger binding than mouse **(Figure S1B)**. These findings support the role of H76, E85, and Q110, with E85Q reducing the affinity of POmAb towards mouse prothrombin.

**Figure 1.**
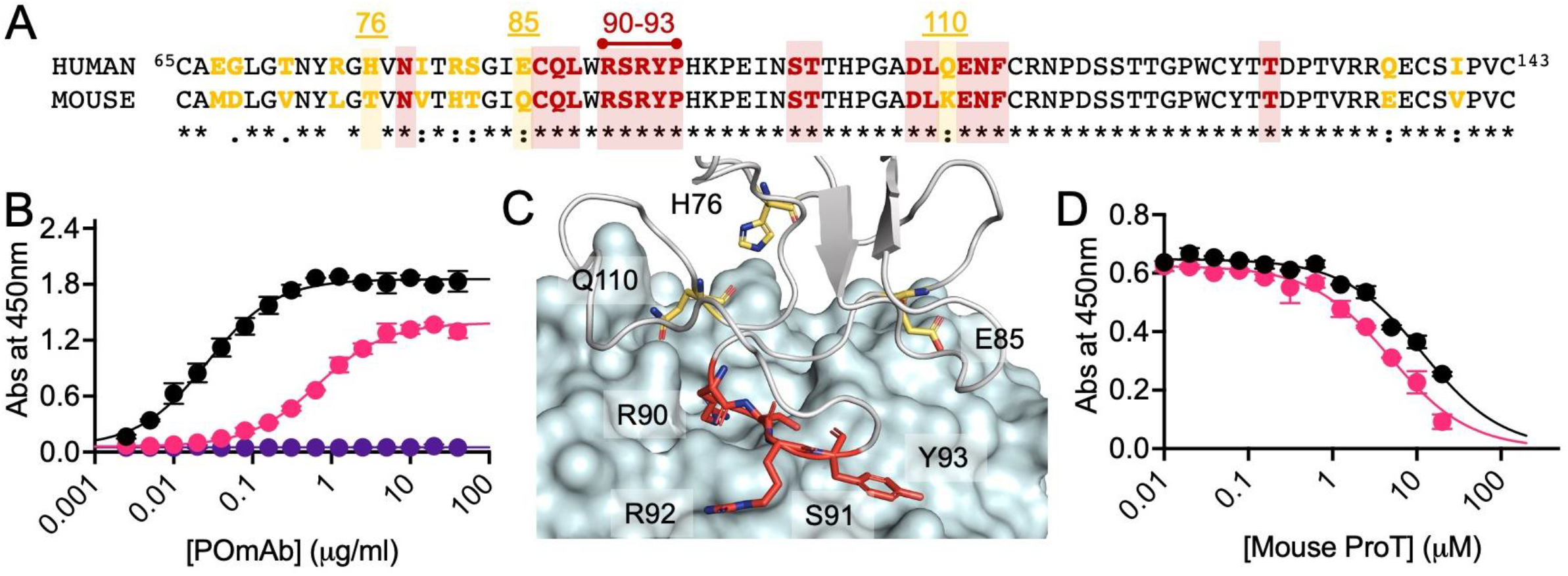
Reactivity of POmAb towards mouse prothrombin. **A)** Sequence alignment of kringle-1 (residues 65-143) of human (P00734) and mouse (P19221) prothrombin. Conserved residues targeted by POmAb, including the region R90-Y93, are highlighted in red, while non-conserved residues are shown in yellow. Residues 76, 85, and 110 are further highlighted as they are integral to the binding interface. **B)** Binding of POmAb to human prothrombin (black dots) and mouse prothrombin (pink dots) measured by ELISA. The negligible binding of the control mouse IgG1 antibody to mouse prothrombin is indicated by purple dots. Best fit of the data was obtained with a Hill equation. EC_50_ values are 0.024±0.005 μg/ml (or 0.16 nM) for human prothrombin and 0.83±0.25 μg/ml (or 5.5 nM) for mouse prothrombin. **C)** Structure of POmAb (cyan surface) bound to kringle-1 of human prothrombin (gray cartoon) shows location of non-conserved residues (yellow stick). The region R90-Y93 (red sticks) is shown for reference. **D)** Zoom-in of the binding interface to highlight the position of residue H76, E85, and Q110 (yellow sticks) that are mutated to T, Q, and K in mouse prothrombin, respectively. **E)** Competition experiments. Inhibition of POmAb binding (10 μg/ml) to immobilized mouse prothrombin (1 μg/well) by mouse prothrombin (0-20 μM, black dots) and mouse prothrombin (0-20 μM) in complex with argatroban (400 μM) (magenta dots). Best fit of the data was obtained with a Hill equation. IC_50_ values are 11.2 ± 1.2 μM for mouse prothrombin and 4.3 ± 0.5 μM for mouse prothrombin in complex with argatroban.

POmAb recognizes an epitope exposed only in the open conformation of prothrombin **(Figure S2A)**. Prothrombin is mainly closed, but argatroban shifts it to the open form^7,8^, enhancing POmAb binding. This effect is consistent in both mouse **(Figure 1E)** and human prothrombin **(Figure S2B),** indicating a conserved conformational mechanism. In competition experiments, the IC_50_ values for mouse prothrombin were 11.2 μM without argatroban and 4.3 μM with it. These are ∼2000- and ∼700-fold lower than ELISA-derived EC_50_ values, suggesting POmAb strongly favors surface-bound over soluble prothrombin, even when prothrombin is already in the open state.

In recalcified mouse plasma, POmAb mildly prolonged the clotting time initiated by dRVV compared to IgG control **(Figure 2A)**. In a chromogenic assay, POmAb reduced the conversion of mouse prothrombin into thrombin by prothrombinase **(Figure 2B)**, although it was less potent than with human prothrombin **(Figure S3)**. These findings confirm that POmAb has an anticoagulant effect in mice *in vitro*.

**Figure 2.**
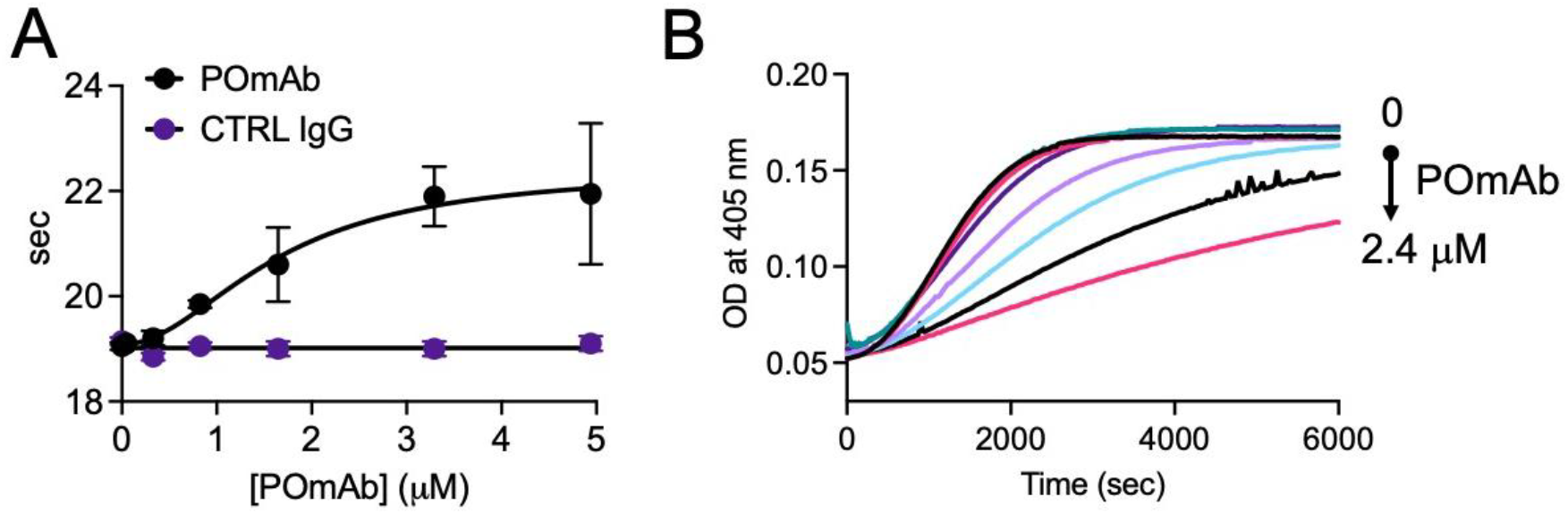
Anticoagulant profile of POmAb *in vitro*. **A)** POmAb-mediated prolongation of the clotting time in recalcified mouse plasma. **B)** POmAb-mediated reduction of prothrombin conversion to thrombin by prothrombinase monitored at 405 nm. The assay began with the addition of a mixture of 2.5 pM human factor Xa, 0.2 nM human Va, and 20 μM phospholipids to 25 nM mouse prothrombin, which had been preincubated for 10 minutes with increasing concentrations of POmAb (0-2.4 μM). Absorbance at 405 nm reports real-time cleavage of the thrombin-specific chromogenic substrate FPFpNA (H-D-Phe-Pro-Phe-p-nitroanilide).

The *in vivo* effect of POmAb was evaluated in a laser-induced cremaster injury model^13-15^, which we previously used to demonstrate that anti-β_2_-glycoprotein I antibodies isolated from APS patients stimulate thrombus formation^17^. Mice received fluorescently labeled POmAb or IgG control (0.5 mg/kg) via the jugular vein. Real-time evolution of thrombus formation was monitored by intravital microscopy using antibodies specific for fibrin (59D8) and platelets (anti-CD42b). Across four mice per group, injury size (40 injuries for POmAb, 48 for control) was comparable **(Figure S4).** Representative images **(Figures 3A, 3B)** and quantification **(Figures 3C–F)** showed that POmAb significantly reduced fibrin formation by ∼3-fold (Ctrl: 77.3 × 10^10^ AFU vs POmAb: 24.1 × 10^10^ AFU; p < 0.001) and had some effect on platelets that however did not reach statistical significance (Ctrl: 2.24 × 10^11^ AFU vs POmAb: 0.85 × 10^11^ AFU; p = 0.1497). POmAb rapidly localized to the injury site **(Figure 3A)**, co-localizing with fibrin and initially with platelets **(Figure 3G)**. The kinetics of prothrombin accumulation are consistent with thrombin being necessary for generating fibrin and activating platelets^18^.

**Figure 3.**
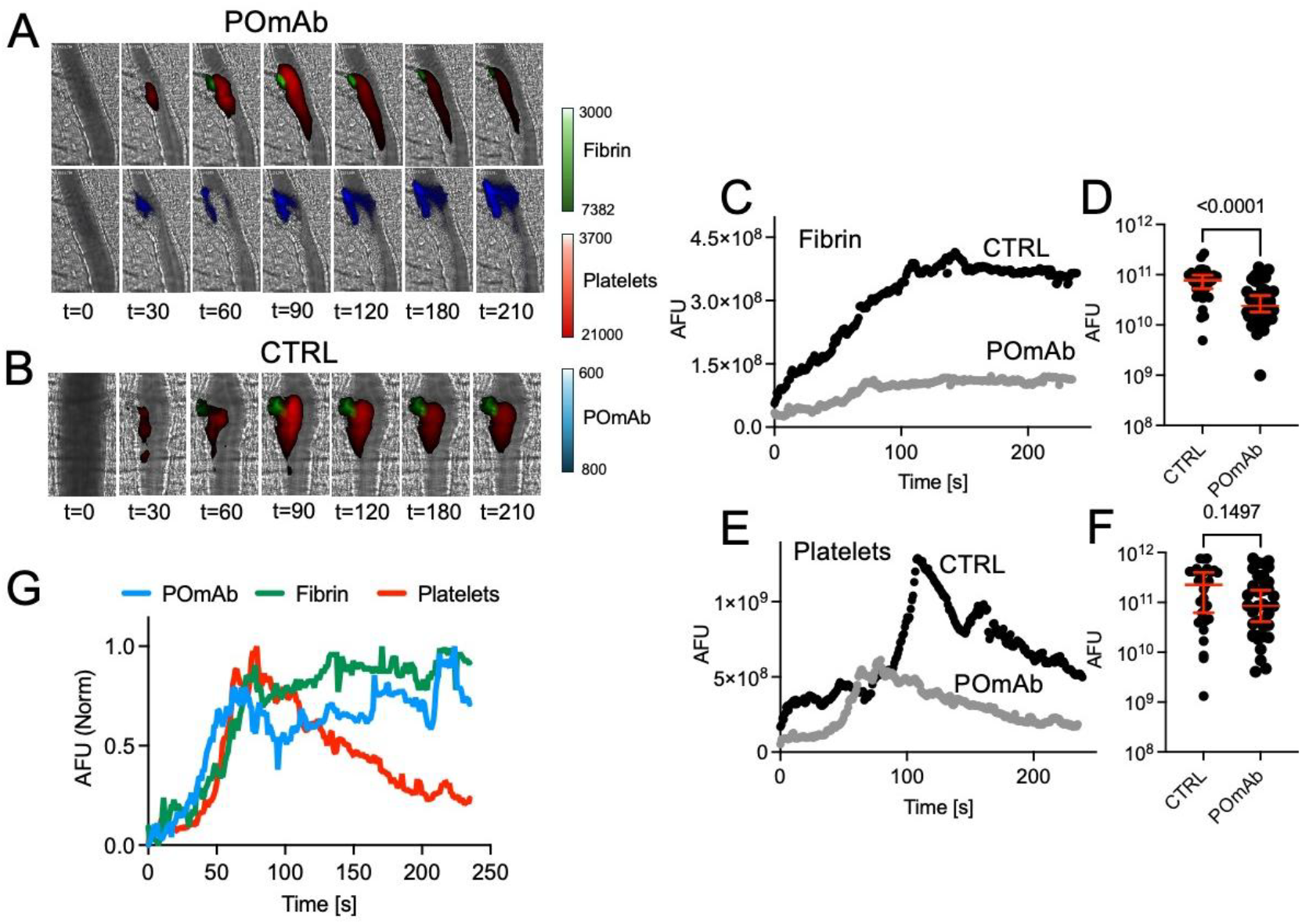
Anticoagulant profile of POmAb *in vivo*. Intravital images of cremaster arterioles were obtained following infusion of **A)** POmAb or **B)** control antibodies. Fibrin is shown in green, platelets in red, and POmAb in blue. **C)** Median integrated fluorescence intensity over time of fibrin is shown. **D)** Quantification of the normalized fibrin accumulation as area under the curve (AUC) with median is shown. CTRL, n = 48 injuries; POmAb, n = 40 injuries. *P*-values were determined by non-parametric one-way ANOVA with Dunnett’s post hoc analysis**. E)** Median integrated fluorescence intensity over time of platelets is shown. **F)** Quantification of the normalized platelet accumulation as area under the curve (AUC) with the median shown. CTRL, n = 48 injuries; POmAb, n = 40 injuries. *P*-values were determined by non-parametric one-way ANOVA with Dunnett’s post hoc analysis. **G)** Median integrated fluorescence intensity over time of DyLight 650-labeled POmAb, Alexa 488-labeled 59D8, and Alexa 405-labeled anti-CD42b are shown.

The substantial reduction of fibrin deposition at the site of injury might predispose animals to bleeding. This was investigated by the tail bleeding assay^16^. Mice received POmAb or IgG control (0.5 mg/kg, via jugular vein), followed by tail transection (1.0 mm). Bleeding time **(Figure 4A)** and total blood loss **(Figure 4B)** were similar between groups (n=11 each; p = 0.6063), with no sex differences, indicating no significant bleeding tendency. POmAb-treated mice showed a higher frequency of re-bleeding events (p = 0.0257; **Figure 4C**). However, this difference did not result in a difference in total blood loss, suggesting a very mild perturbation of hemostasis.

**Figure 4.**
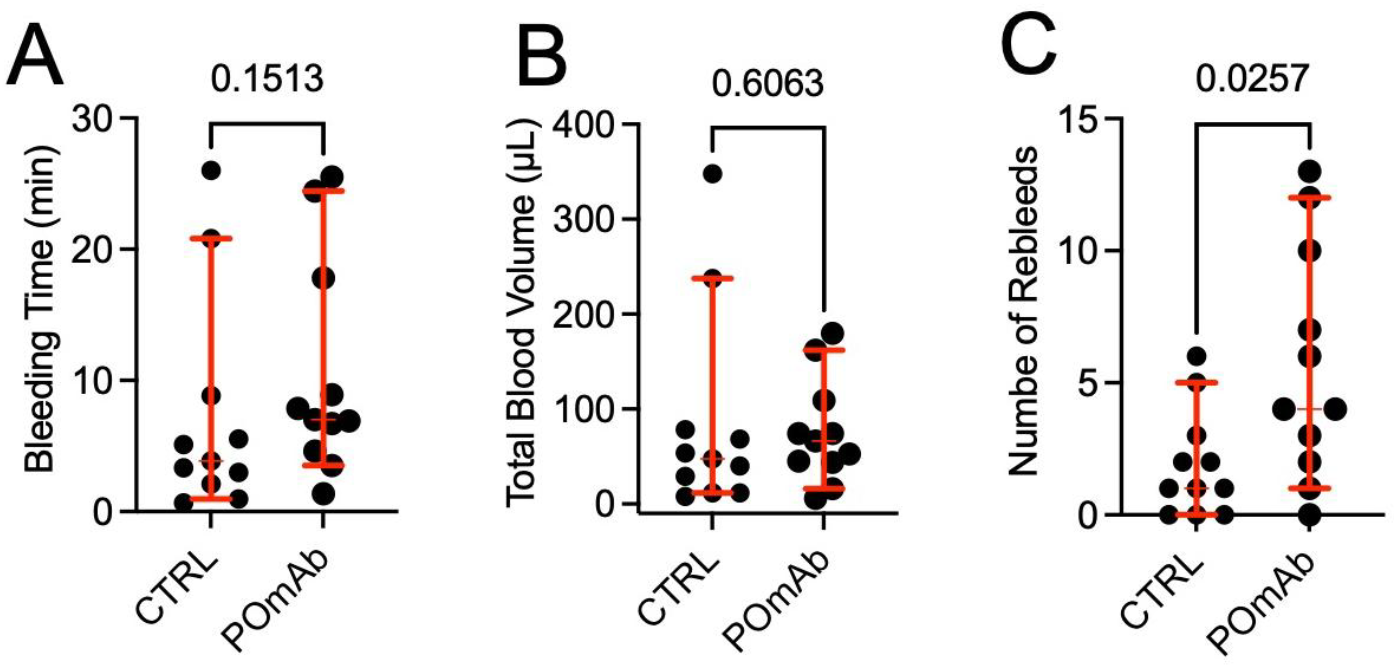
Bleeding propensity. **A-C)** Nine-week-old C57BL/6J mice were evaluated in a tail transection bleeding model and **A)** total bleeding time, **B)** total blood loss, and **C)** number of rebleeds were recorded over 30 minutes of observation for n=11 mice. Mann-Whitney nonparametric test.

## Discussion

POmAb is a prototypical Type-I antibody that targets kringle-1 of prothrombin, keeping it in an open state. Similar to humans, the plasma concentration of prothrombin in mice is estimated to be ∼1.2 μM^19^. In this study, we administered POmAb at 0.5 mg/kg, corresponding to a plasma concentration of ∼55 nM. As an IgG with two Fab arms, POmAb could theoretically bind up to 110 nM prothrombin. The potent anticoagulant effect observed may therefore seem unexpected given the low dose of POmAb compared to prothrombin. However, this outcome is best understood in the context of POmAb’s differential affinities for soluble versus surface-bound prothrombin and its unique mechanism of action.

Equilibrium simulations indicate that, under our experimental conditions, only ∼10% of POmAb binds to circulating prothrombin due to its micromolar affinity **(Figure S5A)**. In contrast, nearly 100% of POmAb is predicted to bind to surface-associated prothrombin, which exhibits nanomolar affinity **(Figure S5B)**. *In vivo,* prothrombin rapidly accumulates at sites of vascular injury^20^, attracted by negatively charged phospholipids exposed on damaged endothelium^21^. This localized environment increases the local concentration of prothrombin and promotes stable POmAb binding to its open conformation mostly through avidity^22-24^. Supporting this mechanism, *in vivo* imaging shows a rapid increase in fluorescence from labeled POmAb at injury sites, along with co-localization with fibrin and, initially, platelets. Importantly, open prothrombin is less efficiently converted to thrombin by prothrombinase^8,10^. Altering prothrombin activation pathway prolongs clotting time in human plasma^12^ and may protect mice against thrombosis^25,26^. Thus, the remarkable anticoagulant effect of POmAb is explained by its selective accumulation at sites of injury and stabilization of the open, less “active” form of prothrombin. How endothelial binding and shear stress enhance POmAb localization remains unknown and warrants further investigation.

Despite the significant effect on fibrin, POmAb had only a modest impact on platelets. This finding is intriguing in light of current understanding of thrombus dynamics in this model. Previous studies using PAR-4 null mice have demonstrated that it is possible to reduce the growth and propagation of platelet thrombi without significantly affecting fibrin formation^27^. However, these mice exhibited markedly prolonged bleeding times in the tail bleeding assay^28^. Studies with POmAb suggest that the opposite result is possible. By targeting prothrombin, POmAb might preferentially affect fibrin formation compared to platelet accumulation, and without causing excessive bleeding. Further studies are needed to elucidate the mechanism underlying this selective effect and to determine its relevance in pathological settings.

While this study demonstrates that POmAb acts as an anticoagulant *in vivo*, it was discovered in the context of APS, a prothrombotic disorder. Previous studies have suggested that anti-prothrombin antibodies play a pathogenic role, including causing larger and more stable thrombi in mice^29^, inhibiting the anticoagulant activity of activated protein C in coagulation tests^30,31^, activating human platelets^32,33^, stimulating thrombin generation^34-36^ and inducing expression of tissue factor and E-selectin on endothelial cells^29,37^, which is unlike what is seen in this study with POmAb. However, there are at least two significant differences to consider. Firstly, most previous studies have either conducted research with mixtures of antibodies purified from APS patients^30,31,35,37^ or with a monoclonal antibody that is polyspecific^29,34,38^. Although POmAb is not derived from humans, it is specific for prothrombin^10^. Second, in this study, POmAb is formatted as a mouse IgG_1_, an isotype that is inefficient at fixing complement^39^. This is in contrast to previously used monoclonal antibodies, which were either human IgM^40^ or human IgG_3_^29^, which efficiently fix complement^41^. Complement hyperactivity is a feature of APS patients^42-44^. Escaping complement fixation by antibody engineering abrogated the pathogenic effect of an anti-domain I antiphospholipid antibody^45^. While more research is needed, this study aligns with the idea that complement plays a significant role in APS^46^ and advances the new provocative concept that not all anti-prothrombin antibodies are necessarily pathogenic; some may be protective, particularly at low concentrations. Unfortunately, mice lack the platelet Fc receptor, FcγRIIa, and this is a limitation in further interpreting these and previous studies in the context of APS.

To summarize, POmAb is a Type-I monoclonal antibody that targets kringle-1 of prothrombin, keeping it in an open conformation. This study demonstrates that, in the cremaster arteriole laser-induced injury model, infusion of POmAb significantly reduces fibrin generation without causing excessive bleeding. Results obtained with POmAb offer new insights into the potential roles of the anti-prothrombin antibodies in APS. They also provide proof of concept for a new class of anticoagulants that, by specifically targeting open prothrombin, could mitigate thrombosis without increasing the risk of bleeding.

## Authorship Contributions

All authors designed the research; M.B., G.M.S., and S.S. performed *in vivo* studies; S.K. produced the antibody and performed *in vitro* studies; all authors analyzed data; N.P., S.S., and R.F. oversaw the project; N.P., S.S., and R.F. wrote the manuscript; all authors edited and reviewed the manuscript.

## Conflict of interest

The authors declare that they have no conflicts of interest with the contents of this article.

## Data availability

All data are contained in the manuscript.

## Funding and additional information

This work was in part supported by grants R01HL150146 (NP), R01HL175060 (SS), R01HL167383 (RF), and R21HL173848 (NP).

## Notes

### Competing Interest Statement

The authors have declared no competing interest.

