## Supplemental Figures for "POmAb, an antibody targeting open prothrombin, results in anticoagulation without excessive bleeding in mice"

A

76 85 90-93 110

```
HUMAN  65CAEGLGTNYRGHVNITRSGIECQLWRSRYPHKPEINSTTHPGADLQENFCRNPDSSTTGPWCYTTDPTVRRQECSIPVC143
MOUSE    CAMDLGVNYLGTVNVTHTGIQCQLWRSRYPHKPEINSTTHPGADLKENFCRNPDSSTTGPWCYTTDPTVRREECSVPVC
BOVINE   CAEGVGMNYRGNVSVTRSGIECQLWRSRYPHKPEINSTTHPGADLRENFCRNPDGSITGPWCYTTSPTLRREECSVPVC
RAT      CAMDLGLNYHGNVSVTHTGIECQLWRSRYPHRPDINSTTHPGADLKENFCRNPDSSTSGPWCYTTDPTVRREECSIPVC
```

B

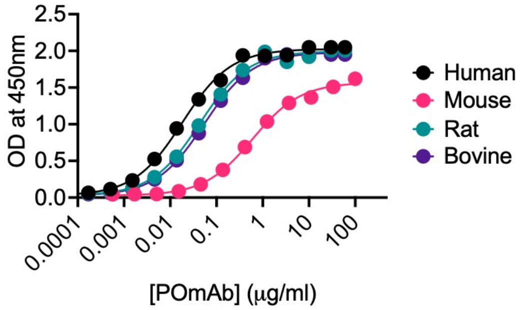


C

| | Human | Mouse | Rat | Bovine |
|---|---|---|---|---|
| Bottom | 0.03 | 0.03 | 0.03 | 0.03 |
| Top | 2.02 | 1.58 | 2.00 | 1.99 |
| IC50 (μg/ml) | 0.018±0.001 | 0.602±0.059 | 0.043±0.003 | 0.054±0.004 |

**Figure S1. Reactivity of POmAb towards human, mouse, rat and bovine prothrombin**. **A)** Sequence alignment of kringle-1 of human, mouse, rat and bovine prothrombin. Residues part of the extended epitope recognized by POmAb are highlighted in red, if conserved and in yellow if not conserved. Note how residue 85 is conserved in human, rat and bovine prothrombin but not in mouse prothrombin. **B)** ELISA assay documenting reactivity of POmAb towards different prothrombin species, which follows the order human > rat = bovine > mouse. **C)** Best fit values obtained with a Hill equation.

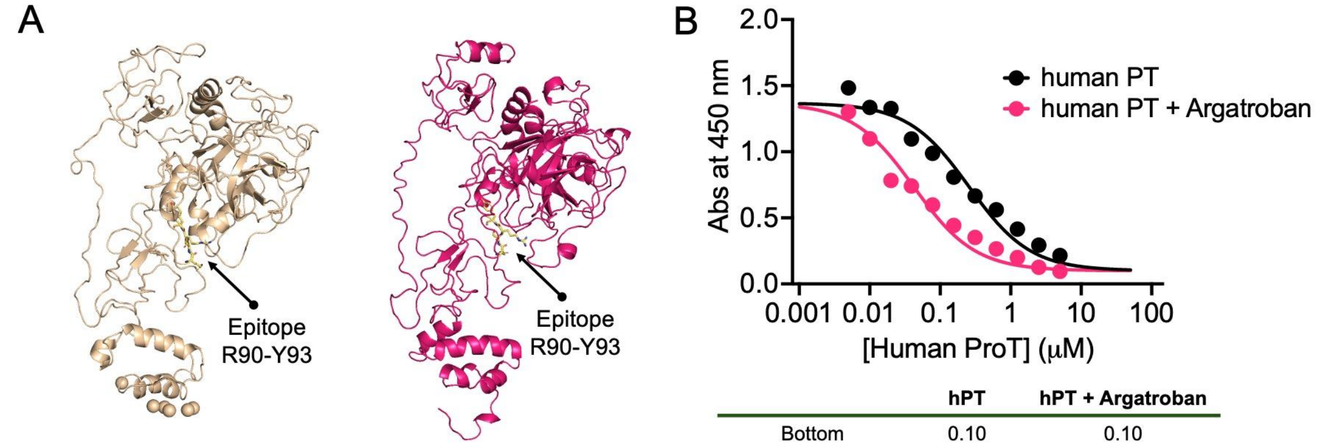


Closed Human Prothrombin (X-ray crystallography – 6CW2)

Closed Mouse Prothrombin (AlphaFold)

| | hPT | hPT + Argatroban |
|---|---|---|
| Bottom | 0.10 | 0.10 |
| Top | 1.43 | 1.43 |
| IC50 (μM) | 0.26±0.05 | 0.042±0.007 |

**Figure S2. Open/closed equilibrium in mouse and human prothrombin**. **A)** Structures of human (wheat) and mouse (pink) prothrombin in the closed conformation showing similar features and location of the epitope recognized by POmAb. Opening of prothrombin is necessary for the epitope to become accessible to the antibody. **B)** Competition experiments. Inhibition of POmAb binding (0.1 μg/ml) to immobilized human prothrombin (1 μg/well) by human prothrombin (0-5 μM, black dots) and human prothrombin (0-5 μM) in complex with argatroban (400 μM) (magenta dots). Best fit of the data was obtained with a Hill equation. $IC_{50}$ values are 0.26 ± 0.05 μM for human prothrombin and 0.042 ± 0.007 μM for human prothrombin in complex with argatroban. Like what is seen in mouse prothrombin **(Figure 1E)**, enhanced reactivity is due to argatroban shifting the equilibrium toward the open form.

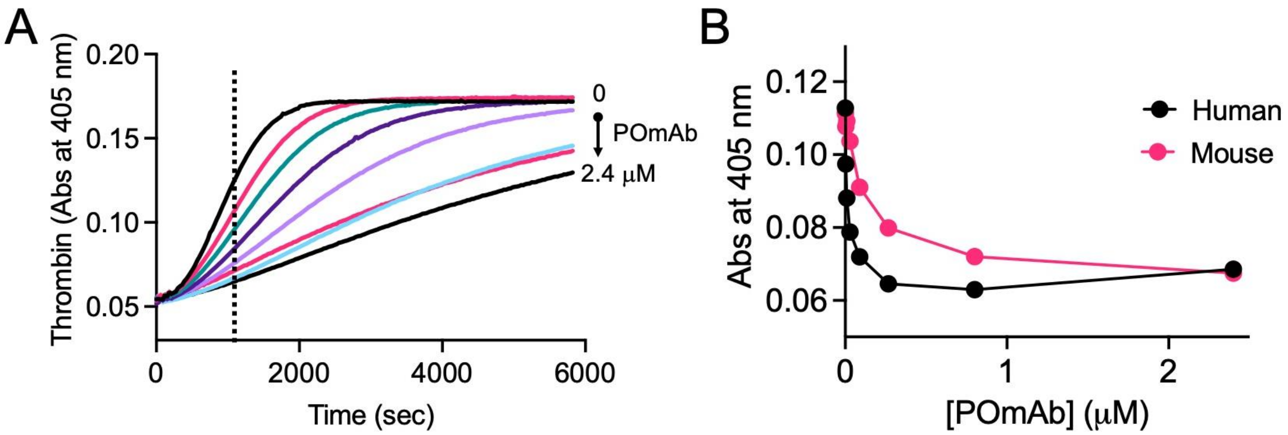


**Figure S3. POmAb-mediated reduction of prothrombin conversion to thrombin by prothrombinase. A)** As in **Figure 1G**, the assay began with the addition of a mixture of 2.5 pM human factor Xa, 0.2 nM human Va, and 20 μM phospholipids to 25 nM human prothrombin, which had been preincubated for 10 minutes with increasing concentrations of POmAb (0-2.4 μM). Absorbance at 405 nm reports real-time cleavage of the thrombin-specific chromogenic substrate FPFpNA. **B)** Absorbance at maximum velocity (960 seconds for human prothrombin and 1230 seconds for mouse prothrombin) plotted against POmAb concentrations, showing that POmAb is ~10-fold more potent against human prothrombin in this assay (0.012 ± 0.016 μM vs 0.123 ± 0.019 μM).

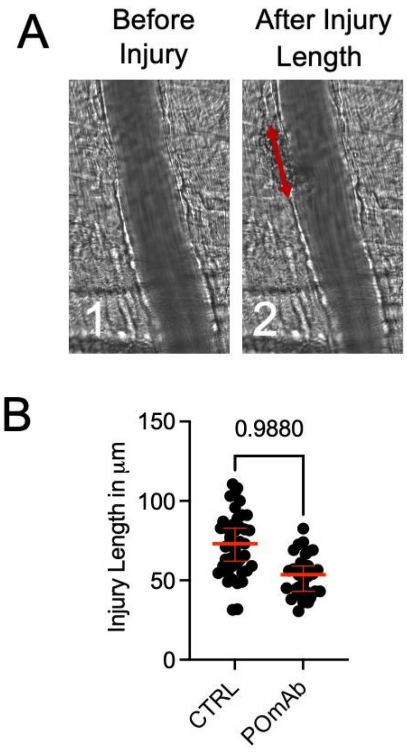


**Figure S4. Injury size**. **A)** Images corresponding to microcirculation before (1) and after the laser injury (2), and criterion used to determine the injury length. **B)** Comparison between control and treated animals. One-way ANOVA test.

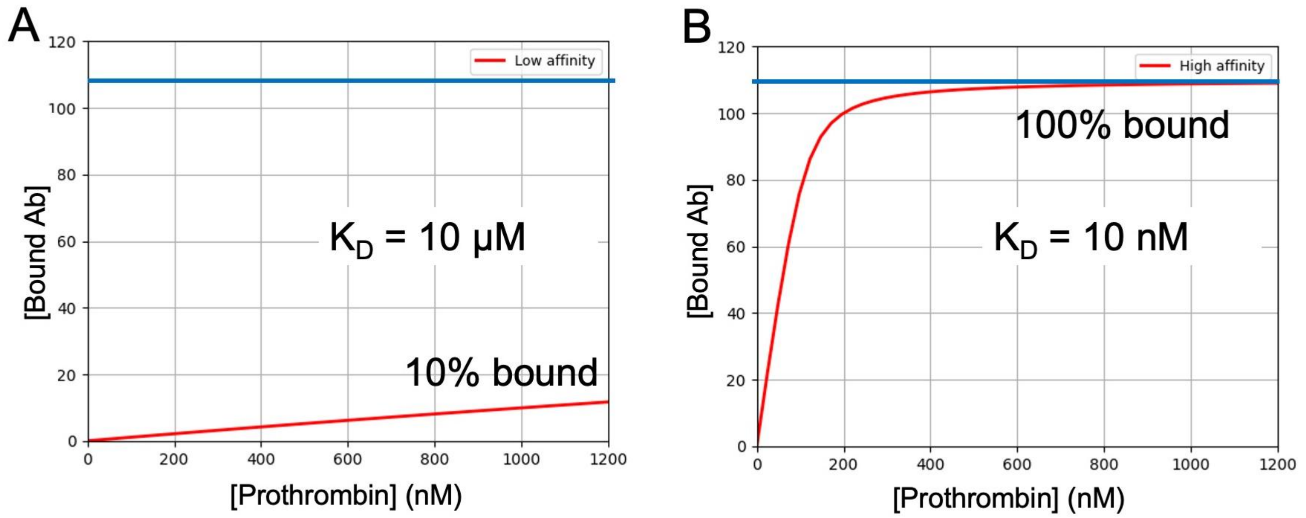


**Figure S5. Equilibrium simulations**. Simulated binding of POmAb (Ab) to prothrombin with an apparent affinity constant of 10 μM **(A,** low affinity**)** and 10 nM **(B,** high affinity**)**, which are estimates determined experimentally. The horizontal blue line indicates a POmAb concentration of 110 nM, representing the maximum binding capacity under our experimental conditions (0.5 mg/kg). At a $K_D$ of 10 nM, 100% of POmAb is expected to bind prothrombin, whereas at $K_D$ = 10 μM, only about 10% (≈12 nM) binds. Given that prothrombin circulates at 1.2 μM (or 1200 nM), this corresponds to approximately 0.1% of circulating prothrombin being bound by POmAb at any given time. These results illustrate why POmAb exhibits minimal binding in the fluid phase and instead rapidly accumulates at sites of injury. Simulations were performed using the Python package PyBindingCurve, as previously described[1].
